# MET functions in tumour progression and therapy resistance are repressed by intronic polyadenylation

**DOI:** 10.1101/2023.08.09.552237

**Authors:** Galina Boldina, Maricarmen Vallejos, Delphine Allard, Mandy Cadix, Céline Labbé, Sophie Vacher, Oskar Hemmingsson, Pierre Gestraud, Aurélie Teissandier, Sylvain Martineau, Didier Auboeuf, Fabrice André, Maud Kamal, Nicolas Servant, Ivan Bièche, Martin Dutertre, Caroline Robert, Stéphan Vagner

## Abstract

Intronic polyadenylation (IPA) leads to the production of transcript isoforms with alternative last exons in thousands of mammalian genes. Widespread regulation of IPA isoforms was observed during oncogenic transformation and in tumours *versus* healthy tissues, and several IPA isoforms were involved in oncogenesis. However, little is known about the potential involvement of IPA in tumour progression, such as cancer cell invasiveness and metastasis, and in resistance to anticancer therapies. Here, we show that an IPA isoform of *MET* (short MET) whose production is inhibited by U1 snRNP (U1), an essential ribonucleoprotein complex that recognizes the 5’ exon-intron junction of pre-mRNA, is associated with better prognosis in breast cancer. Induction of the short MET isoform, using a steric-blocking antisense oligonucleotide targeting the U1 binding site in the vicinity of the short MET alternative polyadenylation site, antagonizes cell invasiveness. U1 blockade with an antisense oligonucleotide targeting the U1 snRNA also decreases breast cancer cell invasiveness, in both human and mouse cancer cell models, and this effect involves IPA induction in *MET* and several genes belonging to the RAS/RAF/MAPK signalling pathway. Finally, short MET relieves melanoma cell resistance to MAPK cascade-targeted therapy *in vitro* and *in vivo*. IPA isoform levels of *MET* and a few other genes (*mTOR*, *EGFR* and *CTNNA1*) help predict such resistance in patients. Altogether, our findings provide evidence for a role of IPA in both cancer cell invasiveness and resistance to therapy. This suggests that IPA isoforms can be exploited as biomarkers and therapeutic targets to combat tumour progression.

## Introduction

The main pathway for 3’ end processing of eukaryotic pre-messenger RNAs (pre-mRNAs) is nuclear endonucleolytic cleavage followed by the addition of a poly(A) tail. This process called cleavage/polyadenylation occurs through the recognition of *cis*-acting elements in the pre-mRNA around the polyadenylation (pA) site by a complex machinery^1^. About 70% of human genes contain alternative polyadenylation (APA) sites^2,3^. The use of APA sites within the same 3’ terminal exon, which is often referred to as tandem APA or 3’UTR-APA, results in mRNA isoforms differing in the length of their 3’ UTR. In contrast, the use of an APA site upstream of the last exon of a gene, which is often referred to as intronic polyadenylation (IPA) not only changes the 3’UTR but also the open reading frame, potentially leading to the production of protein isoforms. Among the many factors that regulate APA, two factors widely repress the expression of IPA *versus* full-length isoforms. The first one is the U1 small ribonucleoprotein particle (hereafter referred to as U1), which represses pA site usage when bound to an upstream 5’ splice site (5’ss)^4,5,6,7,8^. The second factor is cyclin-dependent kinase 12 (CDK12), which promotes transcription processivity and interacts with various splicing factors.

There is now a large body of evidence for a role of 3’UTR-APA in cancer. In particular, 3’UTR shortening is widespread during cell proliferation and in tumours *versus* healthy tissues and has been shown to promote oncogenesis^4,9,10,11,12^. In addition, several studies have linked 3’UTR-APA to tumour progression (*e.g.*, metastasis) and clinical outcome ^13,14^. In contrast, despite early evidence that IPA isoforms are widely regulated in cell proliferation^9^, their widespread regulation in tumours *versus* healthy tissues emerged recently ^15,16^ and only a few IPA isoforms were actively involved in oncogenesis^17^.

Moreover, little is known about the potential role of IPA in tumour progression and clinical outcome. First, there are only two tumour types (*i.e.*, kidney renal clear cell carcinoma and mutiple myeloma) where IPA isoform expression was correlated with patient survival (prognosis)^17,18^. Second, while IPA was involved in cancer cell response to a genotoxic anticancer agent^19^, little is known about the potential role of IPA in the sensibility/resistance to anticancer targeted therapies. Third, whether IPA is widely regulated and functionally implicated in tumour cell invasiveness and metastasis remains to be determined. Indeed, while CDK12 was shown to repress IPA and promote invasiveness in breast cancer cells, a potential link between these two effects was not determined ^20^. Likewise, U1 was shown to regulate HeLa cell migration, but U1 regulated 3’UTR-APA and alternative splicing in these conditions, and therefore no direct link with IPA was demonstrated^21^.

MET, the receptor for HGF (hepatocyte growth factor, or scatter factor SF) is a 190 kDa transmembrane glycoprotein belonging to the family of receptor tyrosine kinases (RTKs) that, upon ligand binding, undergo dimerization and phosphorylate intracellular substrates through their intracellular tyrosine kinase domain, resulting in the activation of the oncogenic MAP-kinase pathway. The HGF-MET axis is activated via diverse events such as DNA amplification, mutations, exon skipping or gene fusions^22^ in multiple cancer types including breast, lung, prostate, head and neck, kidney, pancreatic, digestive, gynecologic, thyroid cancers and melanoma where it is associated with the malignant phenotype and a pejorative prognosis^23–26^. HGF-MET is involved in cancer emergence, progression and resistance to therapies not only as a primary cancer driver event and as a direct drug target, but also as a mechanism of secondary resistance to therapies targeting other factors. For example, it mediates resistance of non-small cell lung cancer to anti-EGFR erlotinib and gefitinib, of HER2 positive breast cancer to anti-HER mAb traztuzumab^27–29^, or of BRAF-mutant melanoma to anti-BRAF vemurafenib therapy^30^. Therefore, targeting MET appears as an attractive therapeutic strategy to overcome anti-EGFR or anti-BRAF resistance^31,32^.

The strong evidence of the involvement of HGF/MET in cancer led to the development of multiple drugs targeting this axis^33^. As of today, only two of them have been approved, both of which are orally available ATP-competitive inhibitors of c-met: Crizotinib, an inhibitor of c-met, ALK and ros-1 authorized for non-small cell lung carcinoma with ROS-1 rearrangement; and cabozantinib for second line treatment of renal cell cancer after failure of anti-VEGFR treatment. Unfortunately, even if these MET inhibitors bring significant benefit to some patients, in most cases resistance eventually occurs^34,35^. Thus, in spite of the huge efforts that have been made for the last 20 years to develop and optimize the use of MET inhibitors in the clinic, it seems that this strategy did not hold its promises with regards to the critical implication of the HGF-MET axis in cancer pathogenesis^36^. In non-small cell lung cancer patients, it appears that the most promising subpopulation of patients prone to respond to MET inhibitors are the ones whose tumours harbor exon-14 skipping whereas tumours with overexpression, amplification or point mutations of MET seem less sensitive^36,37^.

Multiple RTK genes generate IPA isoforms with antagonist properties^8^. In particular, an alternative transcript of *MET* that uses an IPA site in intron 12 encodes a truncated soluble protein isoform (hereafter referred to as short MET) which, when added to cells as an exogenous recombinant protein, was shown to inhibit HGF-induced MET phosphorylation and cell invasiveness^38^. However, the potential roles of the endogenous short MET in cancer have not been investigated. In addition, as stated above, the potential role of IPA in other aspects of tumour progression besides invasiveness, such as resistance to targeted anticancer therapies, is unknown.

In the present study, we show that the endogenous short MET isoform is associated with better prognosis in breast cancer and inhibits tumour cell invasiveness. We also show that U1 inhibition with a steric-blocking antisense oligonucleotide decreases cancer cell invasiveness in both human and mouse, *in vitro* and *in vivo*, and that this effect in human breast cancer cells is dependent on *MET* IPA upregulation. We further show that the large-scale IPA program controlled by U1 in murine mammary tumour cells not only involves MET but also several genes lying in the RAS/RAF/MAPK and PI3K/mTOR signaling pathways downstream of RTKs. Finally, we show that targeted induction of short MET isoform using a steric-blocking antisense oligonucleotide relieves resistance to BRAF-inhibitor therapy in melanoma *in vivo*, and that IPA isoforms of *MET* and other genes can help predict melanoma response to such targeted therapy. Altogether, this study provides strong evidence for a role of IPA in several aspects of tumour progression.

## Results

### The IPA-generated short *MET* isoform correlates with better prognosis and lower invasiveness in breast cancer

To investigate whether the abundance of the IPA-generated *MET* RNA isoform (short MET) relative to the long MET isoform may be associated with tumour progression, we used RT-qPCR with primer pairs designed to specifically amplify the short and long isoforms, as well as all *MET* RNA isoforms (total) (Fig. 1A). We first examined whether the levels of short IPA-produced RNA isoforms were correlated with clinical outcome and could stratify cancer patients based on their prognosis. A Kaplan-Meier univariate analysis of a collection of frozen samples from 416 breast tumour specimens showed a significant positive correlation between low levels of the short *MET* mRNA isoform relative to levels of total (long + short) *MET* (p=0.0065) mRNA and a dismal patient outcome (Fig. 1B), revealing an adverse prognostic value for the IPA-generated *MET* mRNA isoform. Furthermore, in a multivariate analysis using a Cox proportional hazards model, the prognostic significance of the ratio of *MET* mRNA isoforms persisted, indicating that they are independent prognostic factors (p=0.017 and p=0.0049, respectively) (Supplementary Tables S1 and S2). Low levels of short *MET* mRNA isoforms (relative to total *MET* mRNA levels) also predicted poor patient prognosis (p=0.02) in a series of frozen samples from 78 head and neck tumour biopsy samples (Fig. 1C and Supplementary Table S3). Importantly, total *MET* mRNA levels did not correlate with progression-free survival (Fig. 1D). Thus, alterations in short *MET* mRNA levels are associated with progression-free patient survival in two tumour types. Because breast cancer outcome is widely linked to metastasis, we then investigated a potential association of the MET isoform ratio with tumour cell invasiveness. For this, we analysed a panel of 22 human breast cancer cell lines with known strong (n=11) or low (n=11) invasiveness (Supplementary Fig. S1A). This analysis revealed that the ratio of the short to long *MET* mRNA isoforms was significantly lower in cells with a high invasive potential (Supplementary Fig. S1B-S1C). Thus, high levels of the short MET isoform correlate with better prognosis and lower invasiveness in breast cancer.

**Figure 1.**
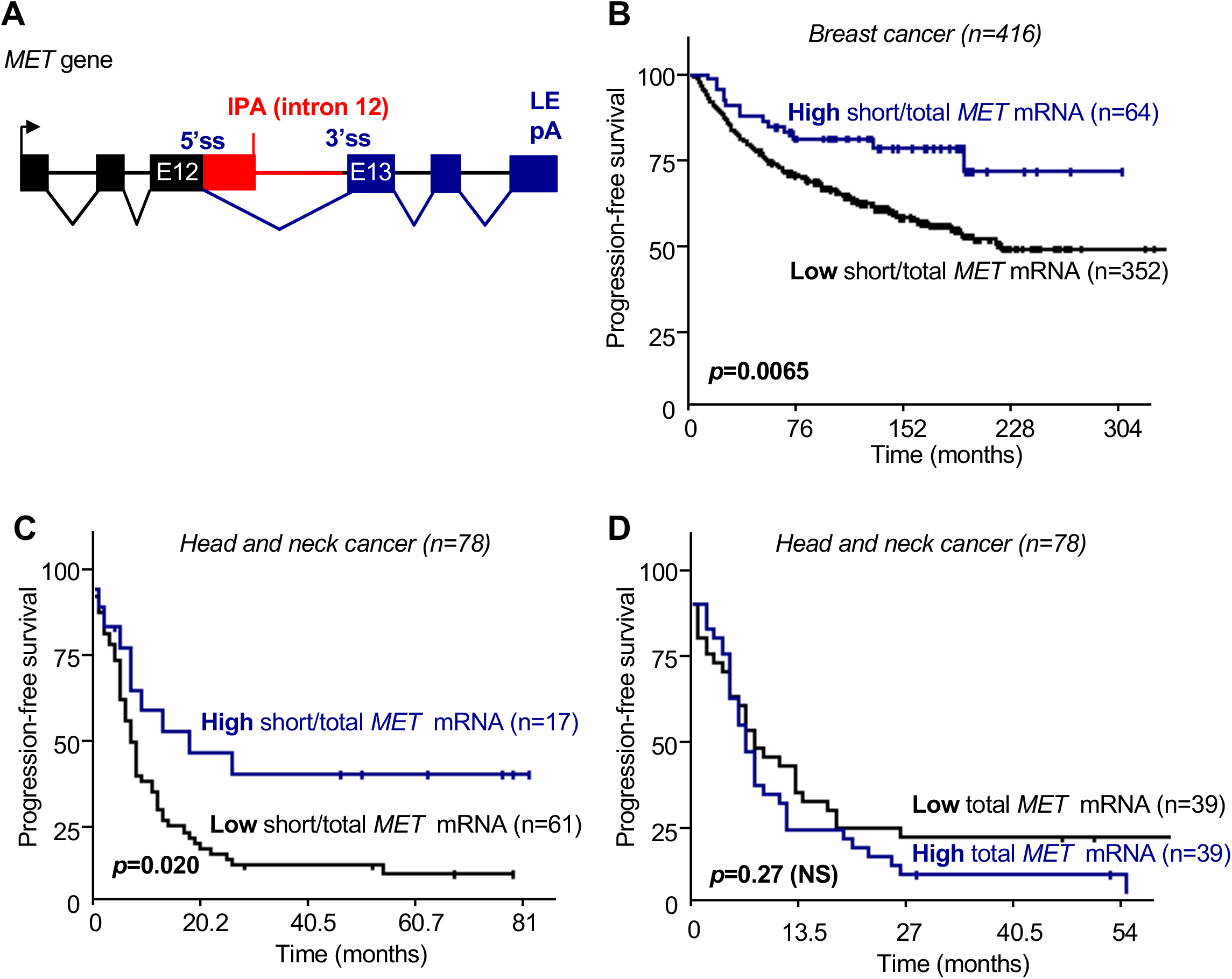
IPA site usage in the *MET* gene is a marker of metastasis-free survival. **A.** Scheme of the *MET* gene depicting the intronic polyadenylation site (pA) located in intron 12 and the last exon (LE) pA. 5’ss: 5’ splice site; 3’ss: 3’ splice site. **B.** Kaplan-Meier analysis of metastasis-free survival (MFS) according to the ratio of short/total *MET* mRNA in 416 breast tumors. **C.** Kaplan-Meier analysis of MFS according to the ratio of short/total *MET* mRNA in 78 head and neck cancer specimens. **D.** Kaplan-Meier analysis of MFS according to the expression level of total MET mRNA in 78 head and neck cancer specimens.

### Targeting of the short *MET* RNA isoform with steric-blocking antisense oligonucleotide regulates tumour cell invasiveness

A previous study showed that an exogenous recombinant short MET protein inhibits cell invasiveness^38^. To assess the role of the endogenous short *MET* RNA isoform in breast cancer cell invasiveness, we transfected an antisense morpholino oligonucleotide (called MET-ASO) that base pairs with the 5’ss of intron 12 of the *MET* pre-mRNA (Fig. 2A). An ASO targeting a given 5’ss will compete with U1 for binding to this 5’ss^8^. In two invasive human breast cancer cell lines (MDA-MB-231 and MDA-MB-468) with low expression of the short *MET* mRNA isoform, MET-ASO led to an increase in the short *MET* mRNA isoforms (relative to the long *MET* mRNA) (Fig. 2B). To ascertain whether this effect was due to regulation of the intron 12 polyadenylation site rather than to a change in the relative stability of the short and long *MET* mRNAs, we measured the ratio of uncleaved (not polyadenylated) to total MET RNA (that is, both cleaved and uncleaved) in nuclear RNA by RT-qPCR^39^ (Fig. 2C). MET-ASO decreased the uncleaved/total RNA ratio indicating that MET-ASO leads to an increased use of the *MET* IPA site (Fig. 2C). MET-ASO treatment also led to increased production of a short MET protein isoform lacking the transmembrane domain and the intracellular tyrosine kinase domain as revealed by western blot with an antibody recognizing the N-terminal part of the protein (Fig. 2D). This short protein isoform was not observed in cells transfected with an siRNA (si-MET (short)) specifically targeting the short isoform (Fig. 2A-2E). We next measured cell migration and invasion *in vitro* in trans-well assays upon HGF stimulation. As expected, HGF stimulated cell migration and invasion (Fig. 2F) as well as increased AKT phosphorylation (Fig. 2G) in both cell lines. The MET-ASO reduced HGF-dependent cell migration and invasion (Fig. 2F) and AKT phosphorylation (Fig. 2G). These data indicate that the endogenous short *MET* isoform counteracts HGF-stimulated MET signaling and tumour cell migration/invasion and demonstrate the feasibility of using an ASO targeting short MET to inhibit breast cancer cell invasiveness.

**Figure 2.**
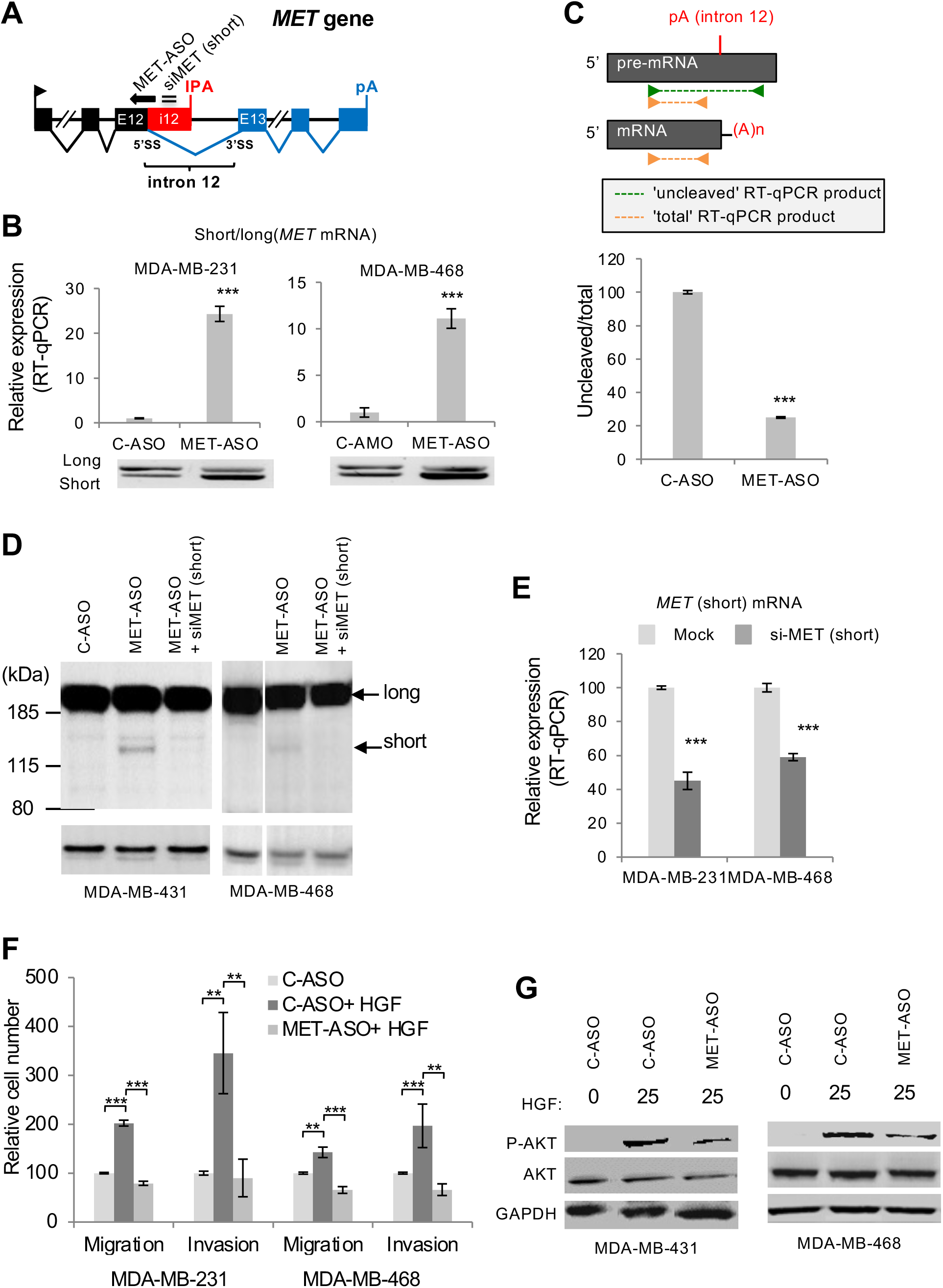
De-repression of *MET* IPA inhibits invasiveness. **A.** Scheme of *MET* pre-mRNA depicting intronic pA (IPA) and pA site within the last exon. The positions of the MET-ASO targeting the 5’ss of intron 12 and of the siRNA targeting the short *MET* RNA isoform (siMET (short)) are indicated. **B.** Expression of *MET* mRNA isoforms (RT-qPCR: upper panel; RT-PCR: bottom panel) in the indicated cell lines treated or not with MET-ASO or a control ASO (C-ASO). Data are presented as the mean ± s.d., and differences were assessed with Student’s t-test (***, P<0.001). **C.** Strategy to evaluate 3’-end processing efficiency at the MET intron 12 polyadenylation site. The primers are located on either side of the cleavage site to amplify the uncleaved RNA, or upstream of the cleavage site to amplify both the uncleaved and cleaved (total) RNA, as depicted. The ratio of uncleaved to total MET RNA is determined by RT-qPCR in nuclear RNA from MDA-MB-231 cells. Data are presented as the mean ± s.d, and differences were assessed with Student’s t-test (***, P<0.001). **D.** Western blot with an antibody recognizing the C-terminal part of the short IPA-generated MET isoform in the indicated cell lines treated or not with MET-ASO and with a combination of MET-ASO and siMET (short). **E.** The efficiency of the siRNA si-MET (short) was tested by RT-qPCR against the short MET RNA isoform. **F.** Migration and invasion assays. HGF: Hepatocyte Growth Factor. Data are presented as the mean ± s.d., and differences were assessed with Student’s t-test (**, P<0.01; ***, P<0.001). **G.** Western blot with antibodies specific for phosphorylated (P-) or total AKT (loading control: GAPDH) in the indicated cell lines treated with MET-ASO or C-ASO.

### U1 blockade decreases human breast cancer cell invasiveness through MET IPA regulation

Because MET IPA was shown to be controlled by U1^8^, we then examined whether U1 could regulate tumour cell invasiveness. To prevent the binding of U1 to 5’ss, we used an antisense morpholino oligonucleotide (called U1-ASO) that targets the 5’ end of the U1 snRNA that base pairs with the 5’ss. ASOs are not substrates for RNase H, and thus do not lead to RNA degradation^8^. Transfection of MDA-MB-231 or MDA-MB-468 with the U1-ASO led to a dose-dependent decrease in cell migration and invasion when compared to a control oligonucleotide (Fig. 3A) and had little effects on cell proliferation (Fig. 3B). Similar effects were observed using siRNA-mediated depletion of the U1-70K protein (a component of U1), but not of the U2AF65 protein recognizing the 3’ss (Supplementary Fig. S2A-S2D), suggesting that the effect of U1 on *in vitro* cell migration and invasion is not merely due to spliceosome inhibition.

**Figure 3.**
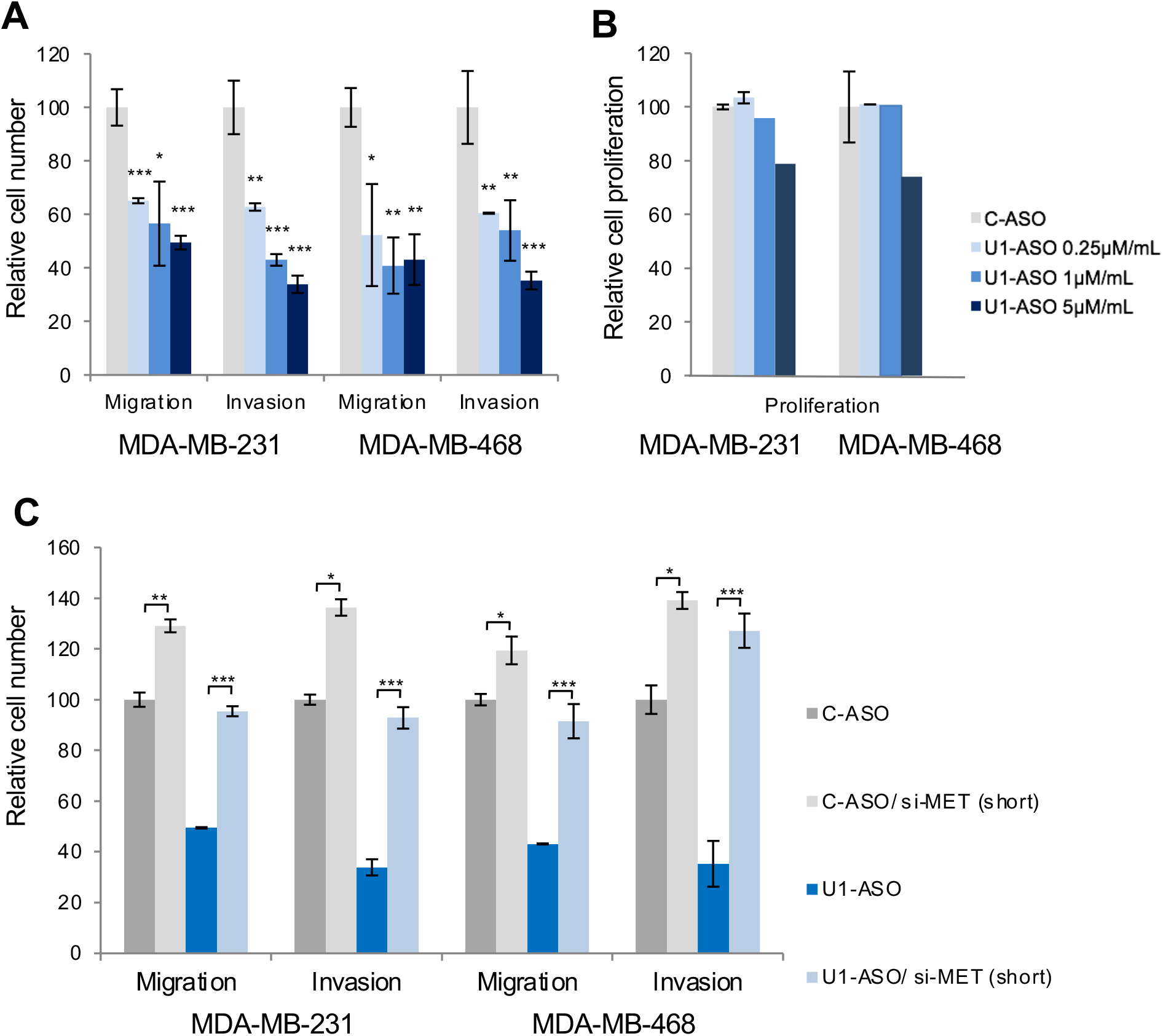
Migration, invasion. (**A**) and **p**roliferation assays (**B**) of MDA-MB-231 and MDA-MB-468 cells after transfection with different concentrations of U1-ASO. Data are presented as the mean ± s.d, and differences were assessed with Student’s t-test (*, P<0.05; **, P<0.01; ***, P<0.001). **C.** Migration and invasion assays of MDA-MB-231 and MDA-MB-468 cells after transfection with the indicated combinations of C-ASO/U1-ASO and si-MET (short) Data are presented as the mean ± s.d, and differences were assessed with Student’s t-test (*, P<0.05; **, P<0.01; ***, P<0.001).

We then evaluated the role of *MET* IPA regulation in U1-dependent regulation of invasive properties. First, both the U1-ASO and the U1-70K siRNA led to an increase in the ratio of short to long MET RNA isoforms in MDA-MB-231 and MDA-MB-468 cells (Supplementary Fig. S2D). Second, depletion of the short MET RNA isoform with the si-MET (short) siRNA described above (Fig. 2D-E) slightly increased the already high migration and invasion capabilities of both MDA-MB-231 and MDA-MB-468 cells in trans-well assays (Fig. 3C), an effect that nicely mirrors the effects of the MET-ASO (Fig. 2F). In addition, the si-MET (short) strongly reduced the inhibitory effect of U1-ASO on both cell migration and invasion in both cell lines (Fig. 3C), suggesting that the effect of U1 in regulating invasiveness is at least in part due to *MET* IPA regulation.

### U1 blockade decreases murine mammary tumour cell invasiveness *in vitro* and *in vivo*

The U1-ASO-dependent decrease in cell migration and invasion in trans-well assays was also observed with the highly invasive 4T1 murine mammary tumour cell line (Fig. 4A), indicating that the role of U1 in these processes is conserved between human and mouse. The U1-ASO also increased 4T1 cell migration as measured by a scratch assay (Fig. 4B). Furthermore, *in vivo* analyses revealed that, following tail vein injection, U1-ASO-treated 4T1 cells were less efficient than control 4T1 cells in developing lung metastasis (Fig. 4C) while primary tumour growth was not affected (Fig. 4D). Altogether, our data indicate that U1 blockade decreases mammary tumour cell migration and invasiveness in both mouse and human, and in various assays *in vitro* and *in vivo*.

**Figure 4.**
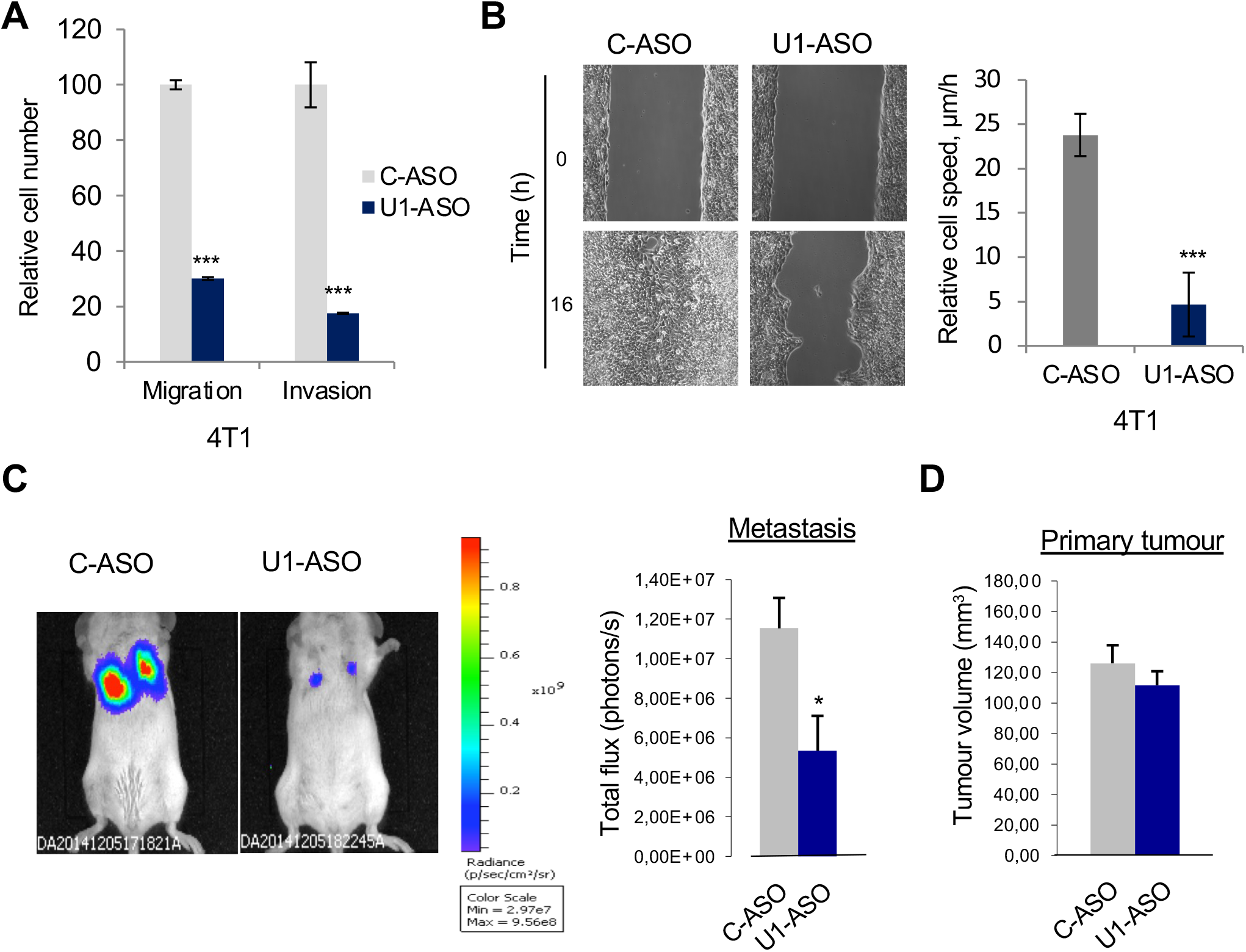
Repression of IPA by U1 promotes invasiveness. **A.** Migration and invasion assays in 4T1 cells after U1-ASO treatment. Data are presented as the mean ± s.d. and differences were assessed with Student’s t-test (***, P<0.001). **B.** Scratch assay after treatment with C-ASO or U1-ASO. Left: representative images. Right: cell migration speed is presented as the mean ± s.d. (n=3), and differences were assessed with Student’s t-test (***, P<0.001). **C.** *In vivo* metastasis assay. U1-ASO-treated versus mock-transfected 4T1-luc2 cells were injected into the tail vein of mice and metastasis to the lung was imaged by bioluminescence (left) and quantitated (right), mean values ± s.e.m.; C-ASO, n=8; U1-ASO, n=7; Mann-Whitney U test; *, P<0.05). **D.** Orthotopic tumor growth assay. Volume of primary tumors 7 days after percutaneous injection in the fourth mammary fat pad of U1-ASO-treated versus mock-transfected 4T1-luc2 cells. Mean values ± s.e.m. are displayed (n=7).

### U1 regulates IPA isoforms in several genes involved in signaling pathways downstream of RTKs

To identify on a genome-wide scale the set of IPA events that are regulated by U1-ASO in 4T1 cells, we used high-throughput sequencing of 3’ ends of polyadenylated transcripts (3’-Seq) (Supplementary Fig. S3A) and our bioinformatics pipeline called 3’-SMART (3’-Seq Mapping Annotation and Regulation Tool) (Supplementary Fig. S3B). We applied this approach to 4T1 cells treated or not with the U1-ASO. The vast majority of sequencing reads were found in the last exon (LE) of coding genes and in the vicinity of a pA signal (AAUAAA), as expected, while about 10% of the reads were found in introns, and intronic pA sites were found in 60% of detected genes (Supplementary Fig. S3D-S3F). 3’-Seq analysis indicated that the U1-ASO led to increased usage of 1113 IPA sites in 841 genes (relative to the sum of pA sites present in the LE of matched genes), and to decreased usage of only 271 IPA sites in 208 genes (p<0.05 and FDR<10%; Fig. 5A and Supplementary Table S4). The majority of regulated IPA sites were annotated in databases of pA sites^40,41^, were significantly expressed (above a threshold of 5% reads relative to the LE) and/or had an AATAAA-like hexamer^42^ (Fig. 5B and Supplementary Fig. S3G-S3H). However, compared to down-regulated IPA sites, up-regulated ones were less often annotated (49% versus 66%), abundant (53% versus 77%) and associated with a canonical hexameric pA signal, AA/TTAAA (29% versus 60%). These data are consistent with previous studies in other cell lines showing that U1-ASO up-regulates IPA isoforms and derepresses cryptic IPA sites in many genes^4,6^.

**Figure 5.**
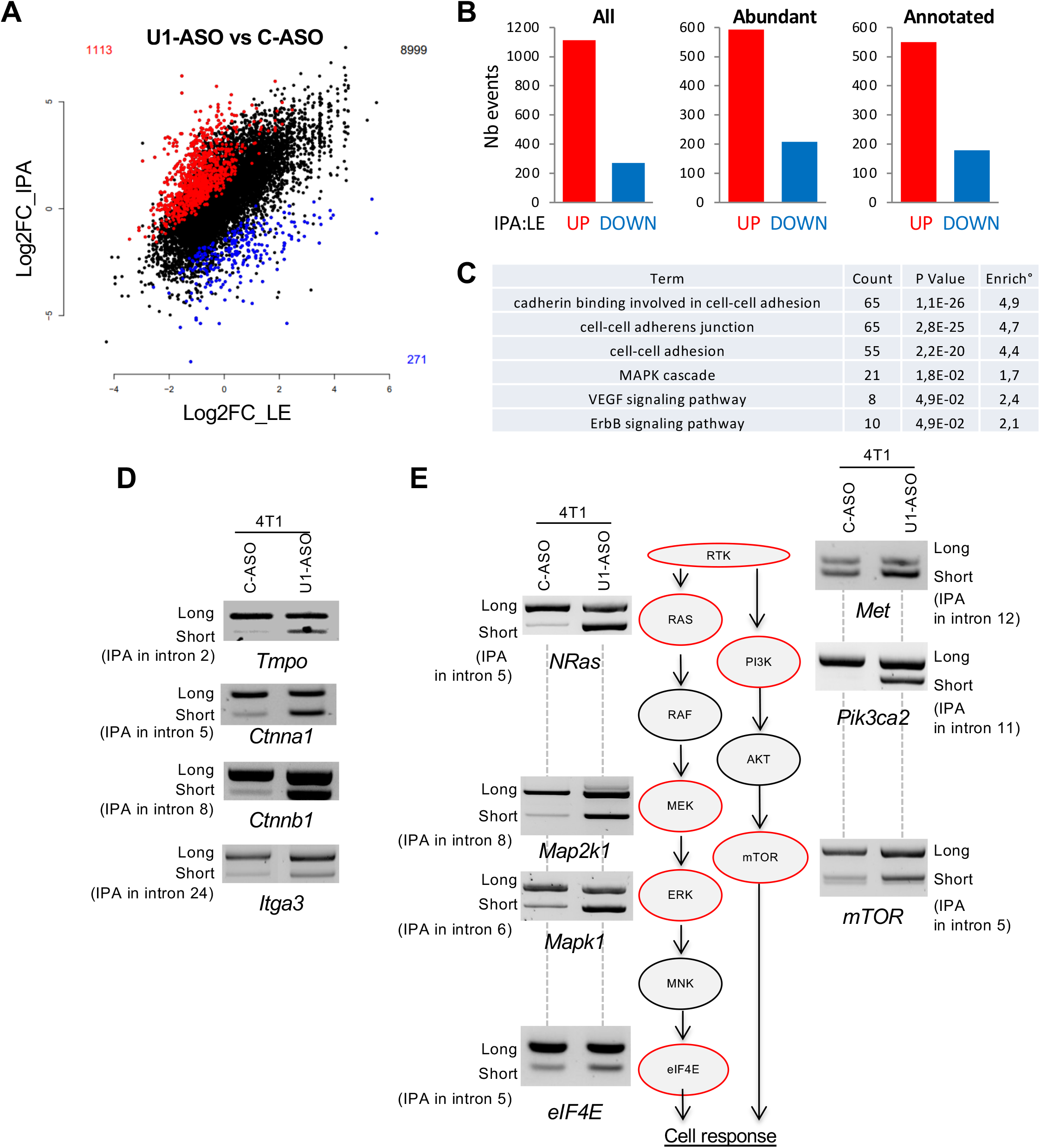
IPA regulation in tumor progression genes. **A.** Regulation of IPA versus LE expression following U1-ASO treatment in 4T1 cells. IPA sites that are significantly increased or decreased by U1-ASO relative to LE (p-adj<0.1) are shown in red and blue, respectively. **B.** Number of genes with increased (red) or decreased (blue) ratio of IPA to LE expression following U1-ASO treatment. **C.** Table showing GO terms enriched for genes with U1-ASO-regulated IPA. **D. E.** RT-PCR analysis showing the relative expression of short and long mRNA isoforms of the indicated genes following U1-ASO treatment in 4T1 cells. RTK, receptor tyrosine kinase.

Interestingly, IPA up-regulation by U1-ASO was enriched in genes involved in specific processes, including cell adhesion and MAPK signaling (Fig. 5C). By RT-PCR, we validated U1-ASO-induced IPA up-regulation events in several genes encoding proteins with functions in cell adhesion and/or migration, including *Tmpo* (a nuclear protein that regulates the motility and metastasis of digestive track cancer cells), *Ctnna1*/α-catenin, *Ctnnb1*/β-catenin and *Itga3*/integrin α3 (Fig. 5D). We also validated IPA up-regulation events in several genes encoding proteins involved in the mitogen activated protein kinase (MAPK) and PI(3)K pathways that lie downstream of RTK signaling, including *NRas, Pik3ca2, Map2k1, Mapk1, mTOR*, and *eIF4E* (Fig. 5E). RT-PCR also showed U1-ASO-induced IPA up-regulation of short *MET* (Fig. 5E), although it was not detected by 3’-seq. Thus, IPA regulation by U1-ASO in 4T1 cells is prominent in signal transduction pathways that are involved in cell migration^43^ and RTK signaling, and are well-known targets of anticancer targeted therapies.

### Short MET antagonizes tumour cell resistance to BRAF inhibitors in melanoma

In addition to promoting tumour cell invasiveness, signaling through HGF-MET has been shown to promote resistance to anticancer therapies. In particular, an important mechanism of innate resistance to RAF inhibitors in melanoma with activating BRAF mutations (*e.g.*, V600E) depends on the tumour micro-environment and more specifically on HGF-MET-dependent reactivation of the MAPK and PI(3)K pathways^30^. We therefore decided to test whether the IPA-generated short MET protein isoform, which lacks the transmembrane domain and the intracellular tyrosine kinase domain and was shown to inhibit HGF-induced MET phosphorylation^38^, could be used to decrease HGF-MET signaling in melanoma cells and to increase their sensitivity to BRAF inhibitors.

Toward these aims, we analyzed MET isoforms expression in 19 melanoma tumour biopsies and 7 melanoma cell lines. In both panels, the isoform ratio was not correlated with total *MET* RNA level (Supplementary Fig. S4A-S4B). MET-ASO treatment of SKMel5 cells, a BRAF-mutated (V600E) melanoma cell line, increased the production of the short MET protein isoform (Fig. 6A) and reduced AKT phosphorylation (Fig. 6B). Similar effects were observed in three other BRAF V600E melanoma cell lines (Fig. 6B) and these results are consistent with our above results in breast cancer cell lines (Fig. 2G).

**Figure 6.**
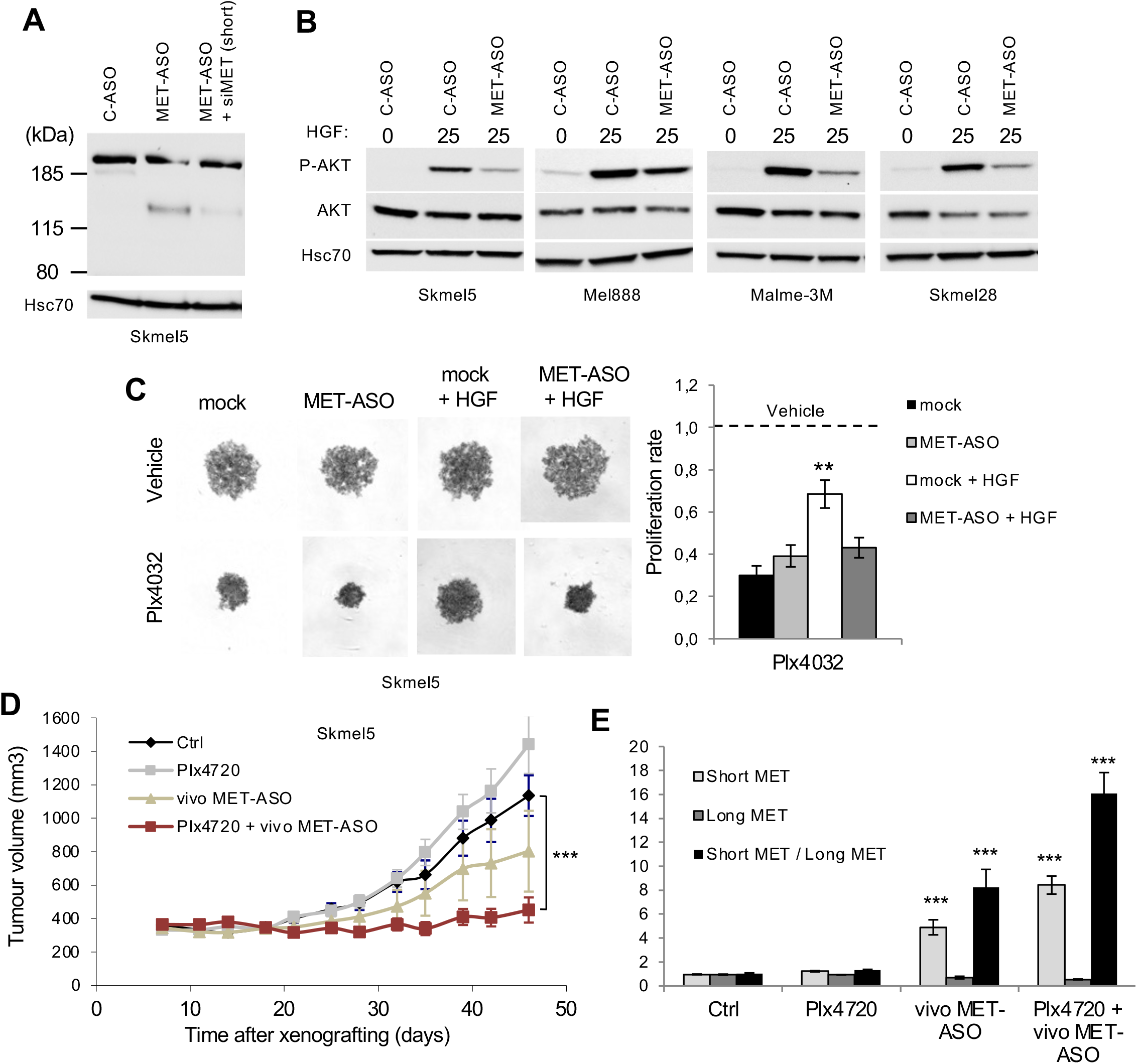
Role of IPA in resistance to anti-cancer therapy. **A.** Western blot with an antibody recognizing the C-terminal part of the short IPA-generated MET isoform in Skmel5 BRAF (V600E) melanoma cells treated or not with MET-ASO and with a combination of MET-ASO and siMET (short). **B.** Western blot with antibodies specific for phosphorylated (P-) or total AKT (loading control: Hsc70) in the indicated cell lines treated or not with MET-ASO. **C.** Spheroids proliferation assay. Representative images of Skmel5 spheroids (on the right) and quantification (on the left) are shown. Histograms represent the mean ± SEM (n=5) and differences were assessed using non-parametric Mann-Whitney U test (**, P<0.01). **D.** Xenograft tumor growth in mice. Curves display the mean ± SEM of 10 tumors per group. Differences were assessed using non-parametric Mann-Whitney U test (***, P<0.001). **E.** RT-qPCR analysis showing the relative expression of short and long MET mRNA isoforms in mouse tumors (mean ± SEM; 10 mice per group); non parametric Mann-Whitney U test; ***, P<0.001).

Then, to investigate the effect of MET-ASO on the sensitivity of SKMel5 cells to BRAF inhibitors, we used two methods. First, we employed a three-dimensional model to grow melanoma spheroids. MET-ASO- or C-ASO-transfected SKMel5 cells were plated as spheroids and treated with a BRAF inhibitor (Plx4032) either alone or combined with HGF. Spheroid growth was not affected by HGF or MET-ASO alone but was inhibited by Plx4032 as expected (Fig. 6C). HGF treatment reduced the inhibitory effect of Plx4032 on spheroid growth, an effect consistent with the study reported by Straussman et al.^30^. Importantly, MET-ASO increased spheroid sensitivity to Plx4032 inhibition (Fig. 6C). This effect was consistent with the fact that MET-ASO decreased HGF-induced MET signaling, as shown above (Fig. 6B). Second, xenograft experiments were performed with MET-ASO covalently linked to a delivery moiety comprising an octa-guanidine dendrimer for *in vivo* purposes (called vivo MET-ASO; see Methods section). The combination of vivo MET-ASO and Plx4032 led to a strong decrease in SKMel5 tumour growth while each treatment given independently had no significant effect (Fig. 6D). This effect on tumour growth was accompanied by a strong increase in the production of the short *MET* RNA isoform (Fig. 6E). Altogether, these data show that MET-ASO treatment can be used to increase the relative levels of the short isoform of MET, decrease HGF-MET signaling, and increase BRAF inhibitor sensitivity in BRAF-mutated melanoma cells.

Finally, we investigated whether IPA isoform levels may be used to predict the response of melanoma tumours to BRAF and MEK inhibitors. To investigate the expression of total and IPA-generated RNA isoforms in clinical specimens, we developed probes for a NanoString nCounter assay (Fig. 7A). This assay was applied to a series of 18 patients who were treated for a metastatic (V600E) BRAF melanoma with BRAF inhibitors (either vemurafenib or dabrafenib) and MEK inhibitors (either trametinib or cobimetinib) (Supplementary Table S5). The biopsies were performed before initiating targeted therapies (baseline samples). The patients were classified as responders, progressors or stable according to Response Evaluation Criteria in Solid Tumours, version 1.1 (RECIST v1.1). Both total and IPA-generated mRNA isoforms of *MET*, but also *EGFR (*Vorlova et al.^8^), *mTOR* and *CTNNA1* (This paper; Fig. 5) mRNAs were evaluated. The expression of total and IPA-generated mRNA isoforms was processed in a factorial discriminant analysis. Total RNA expression only resulted in moderate discrimination between responders, progressors and stable patients (Fig. 7B, left). In contrast, inclusion in the analysis of the IPA-generated short RNA isoforms increased discrimination between the 3 patient categories and raised the percentage of true assignment of patients from 72.22 to 94.44% (Fig. 7B, right). Thus, the quantification of IPA-generated RNA isoforms of several key drivers of cancer increased the possibility of predicting response to treatment.

**Figure 7.**
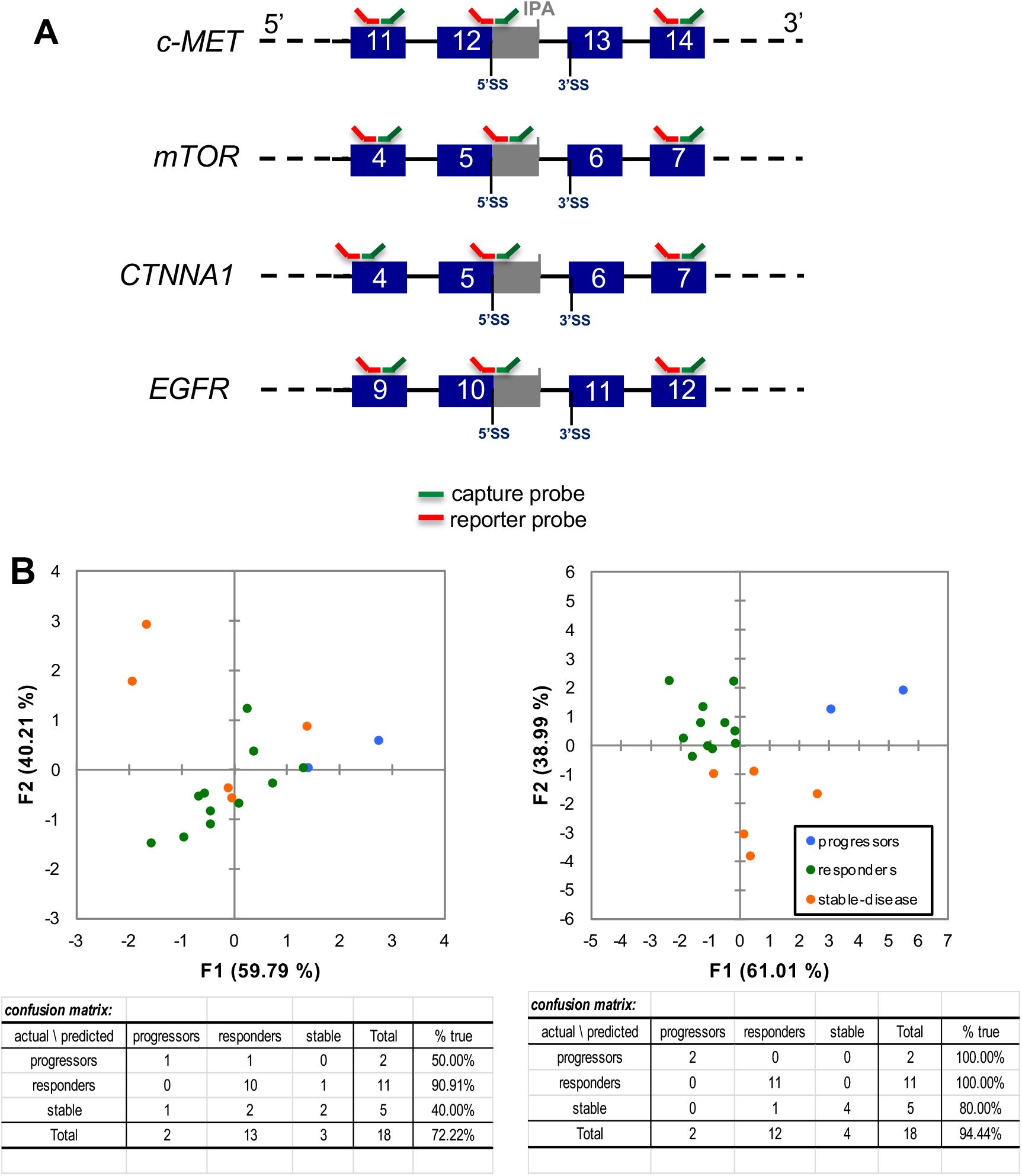
**A.** Schematic representation of capture and reporter probes used in the Nanostring experiments. Capture probes (in green) are complementary to 50 bases of their target messengers and complementary to a Universal Tag which is linked to a biotin used to capture the messengers on a cartridge coated with streptavidin. Reporter probes (in red) are complementary to 50 bases of their target messengers and complementary to a Unique Tag sequence assigned to each target. **B.** Factorial Discriminant Analysis of Nanostring expression data of drivers of resistance to targeted therapies. Total expression (left panel) compared to total + IPA isoforms expression (right panel) in 18 patients under combined anti-BRAF + anti-MEK treatment. Dispersion of patients in two dimensions (up) and confusion matrices (down) are shown.

## Discussion

Metastasis is the overwhelming cause of mortality in patients with solid tumours. Gene expression studies aimed at understanding tumour progression processes and/or at identifying prognostic/theranostic markers often concentrate on steady-state levels of mRNAs or on alternatively spliced variants. Only a few studies have reported individual cases of APA associated with tumour progression^18,45^. We therefore explored the potential impact of APA, and more specifically IPA regulation during tumour cell spreading as well as its influence on response to cancer treatment in various solid tumours.

We show here that the short *MET* RNA isoform generated by IPA is associated with better prognosis and lower invasiveness in breast and head and neck cancers. These findings warrant further investigation regarding the possibility of considering IPA as a predictive marker of treatment. In the specific case of MET and its ligand HGF, a negative correlation was found between the levels of HGF in patient-derived primary melanoma samples and response to therapy in the study by Straussman et al.^30^. However, Lezcano et al.^44^ did not identify a statistically significant correlation. One of the explanations for these discrepancies could be that the level of expression of the short IPA-generated *MET* RNA isoform is an important parameter beyond the raw levels of HGF and MET.

We also report here that targeting of the short *MET* RNA isoform with steric-blocking antisense oligonucleotide regulates tumour cell invasiveness and antagonizes tumour cell resistance to BRAF inhibitors in melanoma. In addition, inhibition of U1 with antisense oligonucleotide decreases human breast cancer cell invasiveness and murine mammary tumour cell invasiveness *in vitro* and *in vivo* and regulates IPA in several genes involved in signaling pathways downstream of RTKs. This highlights the potential therapeutic opportunity of steric-blocking antisense oligonucleotides, especially the one targeting MET, as selective therapeutic tools to be used in combination with well-established anti-cancer targeted therapies. The recent optimization of the use of antisense RNA drugs^46^ makes it possible to rapidly transfer this therapeutic approach to the clinic.

## Materials and Methods

### Cell culture, transfection and treatments

MDA-MB-231 and MDA-MB-468 cell lines (ATCC) were cultured in RPMI-1640 GlutaMax or DMEM:F12 media (Life Technologies) supplemented with 10% FBS (PAN Biotech), 1 mM sodium pyruvate (Life Technologies), and 10 mM HEPES (Life Technologies). 67NR and 4T1 cell lines were kindly provided by F. Miller (Michigan Cancer Foundation, Detroit, Michigan, USA) and cultured in DMEM GlutaMax media (Life Technologies) supplemented with 10% FBS. The A375, mel888, A2058, Skmel28, Skmel5 and Malme-3M melanoma cell lines were purchased from the ATCC. The mel624 cell line was a gift from G. Lizee. Skmel5, Skmel28 and A2058 cells were grown as monolayers in MEM (gibco, #41090) supplemented with 10% heat-inactivated FBS (gibco, #10270); Malme-3M cells were grown as monolayers in IMDM (gibco, #31980) supplemented with 20% heat-inactivated FBS; mel624 were grown as monolayers in RPMI 1640 (gibco, #61870) supplemented with 10% heat-inactivated FBS and mel888 and A375 were grown as monolayers in DMEM (gibco, #31966) supplemented with 10% heat-inactivated FBS. Skmel5 cells were grown as spheroids in DMEM/F12 (gibco, #31331) supplemented with 10% heat-inactivated FBS. All cell lines were cultured at 37°C with 5% CO_2_ atmosphere. Cell lines used were tested negative for mycoplasma contamination using VenorGeM Advance PCR Kit (Biovalley).

Antisense morpholino oligonucleotides (6 µM, unless otherwise stated) coupled to an Endo Porter (6 µl/mL) delivery reagent (Gene Tools, LLC) were added to cell culture media. Transient transfection of siRNAs (50 nM) was performed using Lipofectamine RNAiMAX (Invitrogen, Life Technologies) following the manufacturer’s instructions. All siRNA sequences are provided in Supplementary Table S6.

Ten million of cells were seeded and 24 hours later subjected to C-ASO or MET-ASO transfections. 24h later, cells were stimulated with 50ng/mL recombinant HGF protein (Peprotech) or mock preparation for 15 min.

Vemurafenib (Plx4032) was purchased from Selleckchem (RG7204) and reconstituted as a 10 mM stock solution in DMSO (sigma, D2650). Plx4720 was a kind gift from Gideon Bolag (Plexxikon). Recombinant HGF was purchased from Raybiotech (#228-10702) or Peprotech (#100-39) and reconstituted at 0.5mg/ml in water and then further diluted to a 0.1 mg/ml stock solution in PBS (gibco) 0.1% BSA (Roche, #10735086001). Agarose was purchased from Invitrogen (#16500-500).

### Cell migration and invasion assays

For the cell migration assay, cells (6 × 10^4^ for MDA-MB-468 and 4 × 10^4^ for MDA-MB-231, 4T1) were plated in the upper chamber of an 8 μm pore-size ThinCert (Greiner Bio-One) in serum-free media 24 hours after ASO or siRNA transfection. 10% FBS was added as chemoattractant to bottom chamber. 16 hours later, cells were washed with 1X PBS, fixed with 10% TCA (Sigma) and stained with 1X Amido black (Sigma). Migrating cells were counted from at least three randomized fields per well under an inverted microscope (Leica DM IRBE). The cell invasion assay was performed in similar conditions, except that wells were coated with 100 µl of Matrigel (BD Biosciences) at 1:6 dilution for 4T1 cells or 1:10 dilution for MDA-MB-231 and MDA-MB-468 cells. In the figures, the histogram bars represent the number of migrating/invasive cells as a percentage of the control sample. Each assay was done in duplicates and repeated independently at least three times.

### Scratch assay

16 hours after ASO or siRNA transfection, 8 x 10^3^ cells were plated into chambers separated by silicon insert (culture-insert in µ-Dish 35 mm, Ibidi) allowing formation of confluent monolayer. 24 hours later, the culture medium was replaced with a serum-free medium in order to impair cell proliferation and a wound was made by removing the insert from the plate. Cell migration was followed by live imaging over 16 hours at 37°C with 5% CO_2_ atmosphere. Pictures were taken every 30 min using LEICA DMI 6000B microscope (x10 magnification). The cell migration speed was then calculated based on the 0 and 16 hours time points.

### Western blot analysis

Western blot analysis was performed on cell lysates prepared using RIPA buffer (Sigma). Proteins were quantified using the Pierce™ BCA Protein Assay (Thermo Scientific, # 23225). Samples were resolved in NuPAGE® 4-12% Bis-Tris gels using MES buffer (Invitrogen, Life Technologies, # NP0321) and transferred to a 0.45 µM nitrocellulose membrane (GE Healthcare). After saturation in Tris-buffered saline buffer supplemented with 5% powdered milk, membranes were incubated with antibodies (diluted at 1:1,000) overnight at 4°C. Antibodies specific for the following proteins were purchased from Cell Signaling Technology: MET (mouse, 3148S), MET phospho-Tyr1234/1235 (rabbit, 3077S), AKT (rabbit, 9272), AKT phospho-S473 (rabbit, clone D9E, 4060), ERK1/2 (rabbit, 9102) and ERK1/2 phospho-T202/Y204 (rabbit, clone 20G11, 4376). The antibodies specific for N-terminal MET (Rabbit, ab51067) and U1-70K (Rabbit, ab83306) were purchased from Abcam. The primary antibody for Hsc-70 (sc-7298) was purchased from Santa Cruz. The GAPDH monoclonal antibody (Mouse, G8795) was purchased from Sigma. Horseradish peroxidase (HRP)-conjugated secondary antibodies were purchased from Sigma. In most cases the displayed images were cropped to focus on the protein of interest.

### RNA extraction and RT-PCR

Total RNA was extracted using TRIzol reagent (Invitrogen, Life Technologies, # 15596026). After DNAse I treatment (TURBO DNA-free, Ambion, # AM1907) the RNA was quantified using a NanoDrop 2000 instrument (Thermo Scientific) and reverse transcribed using SuperScript III and random hexamers (Invitrogen, Life Technologies) according to the manufacturer’s instructions. PCR reactions were performed using 10 ng of cDNAs (RNA equivalent) and the GoTaq DNA polymerase (Promega, # M8295). For each gene, a common forward primer and two reverse primers, specific to a full-length or a short form of mRNA, were used. All primers sequences are listed in Supplementary Table S7. Quantitative PCR (qPCR) reactions were performed using 5 ng of cDNAs (RNA equivalent) and the Power SYBR Green Master mix (Applied Biosystems). Data was analyzed by the ΔCt methods and normalized to TBP expression.

### 3D culturing

Skmel5 cells, transfected for 24 hours as monolayers with MET-ASO and Met short siRNA2 or siCtrl sequence, were trypsinized (Gibco, #25300) and plated in round-bottomed 96-well plates coated with 1% agarose in water at the density of 2 x 10^3^ cells per well in DMEM/F12 + 10% SVF medium. Three hours later, cells were treated with HGF and Plx4032 at 50 ng/ml and 1 µM final concentration respectively. Cells were then maintained for 48 hours at 37°C 5% CO2 in a humidified incubator. At the end of this incubation time, the proliferation of spheroids was measured using WST1 (Roche) following the manufacturer’s instructions. Spheroids were mechanically dispersed by pipetting up and down with a multichannel pipettor right after adding the WST1 compound and before a 2-hour-incubation time at 37°C.

### 3’-Seq library preparation and sequencing

5 µg of total RNA was used for poly(A)+ RNA purification with MicroPoly(A)Purist™ Kit (Ambion). Poly(A)+ RNA was fragmented at 70°C for 10 min in RNA Fragmentation buffer (Ambion). After ethanol precipitation, RNA was reverse transcribed using anchored oligo-dT and SMART Scribe RT enzyme (Clontech) allowing to add adapter sequences to the 5’ and 3’ ends of RNA fragments. Libraries were amplified by 13 cycles of PCR with Phusion polymerase (New England Biolabs). A size selection step was done using Agencourt AMPure XP (Beckman Coulter) magnetic beads to obtain fragments of 200–300 bp. Purified libraries were subjected to sequencing using the HiSeq 2000 machine (Illumina). 50 bp reads were obtained by single-end sequencing with P5 Illumina primers. RT and PCR primers are listed in Supplementary Table S7.

### 3’-Seq analysis

Raw reads were trimmed in their 5’ and 3’ ends to remove uninformative nucleotides due to primer sequences, nucleotides added by SMART Scribe RT enzyme and polyA tail of mRNAs. Trimmed reads of 25 bp or more were aligned on the Mouse reference genome (mm9) using Bowtie2 (version 2.2.5) (Langmead and Salzberg, 2012). Only reads with a mapping quality score (MAPQ) of 20 or more were retained (Samtools version 1.1) for downstream analysis (Table S8). Reads were then clustered along the genome using Bedtools (version 2.17.0) (Quinlan and Hall, 2010) allowing a maximum distance of 170 bp (max library fragment length (300bp) – min fragment length (60bp) – read length (50bp) – oligodT length (25bp)) and a minimum number of 5 reads per peak. Peaks with a stretch of 6 consecutive As (or 8 As out of 9 nucleotides) within 150 bp downstream were filtered out, as they are likely due to internal priming of oligo-dT. Overlapping peaks from all analyzed samples were merged to define a common set of genomic windows corresponding to polyA sites.

To annotate peak location within genes, gene coordinates were obtained on the basis of overlapping Refseq transcripts with the same gene symbol. Peaks overlapping any intronic region of a gene were classified as intronic polyA (IPA) peaks. Peaks overlapping the last exon of a gene were classified as LE peaks. Differential analyses between two conditions were done using two independent biological replicates per condition. To compare the regulation of each IPA to the regulation of the gene’s last exon (taken as the sum of the peaks in this exon), we used DESeq2 (version 1.4.5) (Love et al., 2014) and the following statistical model:

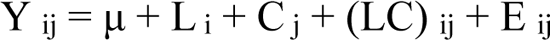

where *Y _ij_* is the normalized counts of peak i in biological condition j, μ is the mean, *L _i_* is the peak localization (IPA or LE), *C _j_* is the biological condition, (LC)_ij_ is the interaction between peak localization and biological condition, and E_ij_ is the residual. Adjusted *P*-values (Benjamini-Hochberg) were calculated. The complete bioinformatics pipeline (3’-SMART package) described above and in Supplementary Fig. S3B can be freely downloaded at GitHub (https://github.com/InstitutCurie/3-SMART) and can be run through a configuration file and a simple command line.

### Clinical samples, quantitative PCR and statistical analysis

Total RNA was extracted from tumour samples by using acid-phenol guanidium. RNA quality was determined by electrophoresis through agarose gels, staining with ethidium bromide and visualization of the 18S and 28S RNA bands under ultraviolet light. RT-qPCR was performed with the ABI Prism 7900 sequence detection system (Applied Biosystems) and PE biosystems analysis software according to the manufacturer’s manuals. Results, expressed as N-fold differences in target gene expression relative to the *TBP* gene and termed “N*target*”, were determined as N*target* = 2^11Ct*sample*^, where the ΔCt value of the sample was determined by subtracting the average Ct value of the target gene from the average Ct value of the *TBP* gene. Primer sequences are listed in Table S7. Agarose gel electrophoresis was used to verify the specificity of PCR amplicons. To visualize the efficacy of a molecular marker (mRNA level) to discriminate between two populations (patients that developed/did not develop metastases) in the absence of an arbitrary cut-off value, data were summarized in a ROC (receiver operating characteristic) curve (Hanley and McNeil, 1982).

### Orthotopic tumour growth and metastasis assays

4T1-luc2 (Caliper Life Sciences) cells, which stably express the Firefly luciferase gene, were transfected with U1-AMO or mock preparation as described above. 24 hours after transfection, cells were trypsinized, resuspended in DBPS and counted. For orthotopic tumour growth assays, 3.5 x 10^5^ cells were injected percutaneously into the fourth mammary fat pad of mice. For metastasis assays, 1 x 10^5^ cells were injected in the tail vein of mice. Six- to eight-week-old female BalbC/Cj mice were used (Janvier, France). Orthotopic tumour growth at the site of injection was assessed 7 days after injection by caliper measurements in two dimensions; tumour volumes were calculated as the volume of an ellipsoid using the formula V(mm^3^) = L(mm) x w(mm)^2 x 0,52 (L=Length, w=width). Metastatic lung colonization was monitored by bioluminescence imaging (IVIS50) 7 days after intravenous injection. Luciferin (K salt, Promega) was injected intraperitoneally at a dose of 150 mg/kg of body weight 15 minutes before imaging. No randomization was used to allocate mice to the different treatment groups and the investigator was not blinded to the group allocation during the experiment. In the orthotopic graft protocol, 7 mice treated with U1-ASO were compared to 7 mice treated with C-ASO. In the metastasis assay, 7 mice treated with U1-ASO were compared to 8 mice treated with C-ASO. Bioluminescence imaging and animal procedures were conducted in the PFIC and PFEP facilities of the Gustave Roussy Institute, respectively, and were approved by the local ethics committee for animal experimentation (CEEA N°26 registered by the Ministère de la Recherche; approved project number Apafis#503).

### nCounter Gene Expression analysis

Total RNAs of patients were extracted from either FFPE (Formalin-fixed paraffin-embedded) or frozen sections (5 x 20µ) using RecoverAll Total Nucleic Acid Isolation (Ambion) or Trizol (Ambion) respectively. Contaminant genomic DNA was removed by a DNase I digestion step using TURBO DNA free kit (Ambion). Tumour cells percentages were evaluated on HES sections by a pathologist (G. Tomasic) and samples showing less than 50% tumour cells were excluded. DNA-free RNAs were quantified with Nanodrop 2000 (Thermo Scientific) and qualified with Bioanalyzer 2100 (Agilent Technologies).

Custom codesets specific for short polyadenylated, full length and total forms of c-MET, mTOR, CTNNA1 and EGFR were designed by Nanostring. ACTB, GAPDH and TBP were used as housekeeping genes. Targeted sequences are given in Table S9. Reporter codesets counts were corrected with the internal negative controls and normalized to positive controls and housekeeping genes. Normalized results were analyzed using the factorial discriminant analysis statistical program in the XLSTAT software.

### *In vivo* experiments

Five-to-eight weeks-old female athymic nude (nu/nu) mice (Gustave Roussy breeding, Villejuif) were engrafted with 5 x 10^6^ Skmel5 cells in 200 µl of DPBS-50% Matrigel (BD Biosciences, #354234) on the right flank. When the tumours were established, mice were separated in 4 groups of 10 mice with comparable mean tumour volumes. No randomization was used to allocate the mice to the different treatment groups. Twenty mice (two groups) were fed with Plx4720-diet (90 mg/kg, Research diets) ad libitum either alone (n=10) or in combination with vivo MET-ASO (125 µM, n=10) injected intratumourally twice a week (as indicated by black arrows on graph B). The 10 mice treated by Plx4720-diet alone received intratumoural (IT) injections of PBS. We evaluated that for a mouse of 22 grams eating 3,5 grams of diet a day the treatment dose of Plx4720 is about 14 mg/kg/day. Twenty mice were fed with the control chow diet (Research diets) ad libitum. Ten of them received IT injections of vivo MET-ASO twice a week. Tumour dimensions were measured with a caliper in two dimensions (L=Length; l=width) twice a week and tumour volumes were calculated using the formula L x W^2 x 0.52. All procedures were approved by the local ethics committee for animal experimentation (CEEA N°26 registered by the Ministère de la Recherche; approved project number Apafis#547).

## Data Availability

All data generated in this study are available upon request from the corresponding author.

## Acknowledgements

This research was funded by the *CNRS*, Curie Institute and *Ligue Nationale contre le Cancer* (Equipe labellisée) to SV. High-throughput sequencing was supported by the grant ANR-10-EQPX-03 from the *Agence Nationale de la Recherche* (*investissements d’avenir*). We thank the Institut Curie Next Generation Sequencing (Thomas Rio-Frio and Sylvain Baulande) and Genomics (Audrey Rapinat and David Gentien) platforms for high-throughput sequencing and Nanostring measurements, respectively. Acquisition of the Nanostring platform was initiated and supported by ANR-10-IDEX-0001-02 PSL, ANR-11-LBX-0044, and Grant « INCa-DGOS-4654 » SIRIC11-002. MV was supported by a post-doctoral fellowship from the “*Association pour la Recherche sur le Cancer*” (ARC, n°PDF20140600985). MC was supported by the grant FRM-DBI20141231314 from the *Fondation pour la Recherche Médicale* to MD. OH was supported by post-doctoral fellowships from the Wenner-Gren foundation and the Swedish Society of Medicine.

## Author contributions

SVag, CR, GB, MV and DA conceived the project. GB, MV, DA and OH conceived and performed experiments. SVac and IB performed the RTqPCR experiments on patient samples. The bioinformatics/biostatistics analysis were supervised by MD and NS and were performed by GB, MC, CL, PG and AT. DA, FA and MK provided clinical samples and/or gave advices. CR supervised the work on melanoma. SVag supervised all research and wrote the manuscript with critical input from GB, MV, DA, IB, CR and MD. All other authors edited the manuscript.

---

**Supplementary Figure 1.**
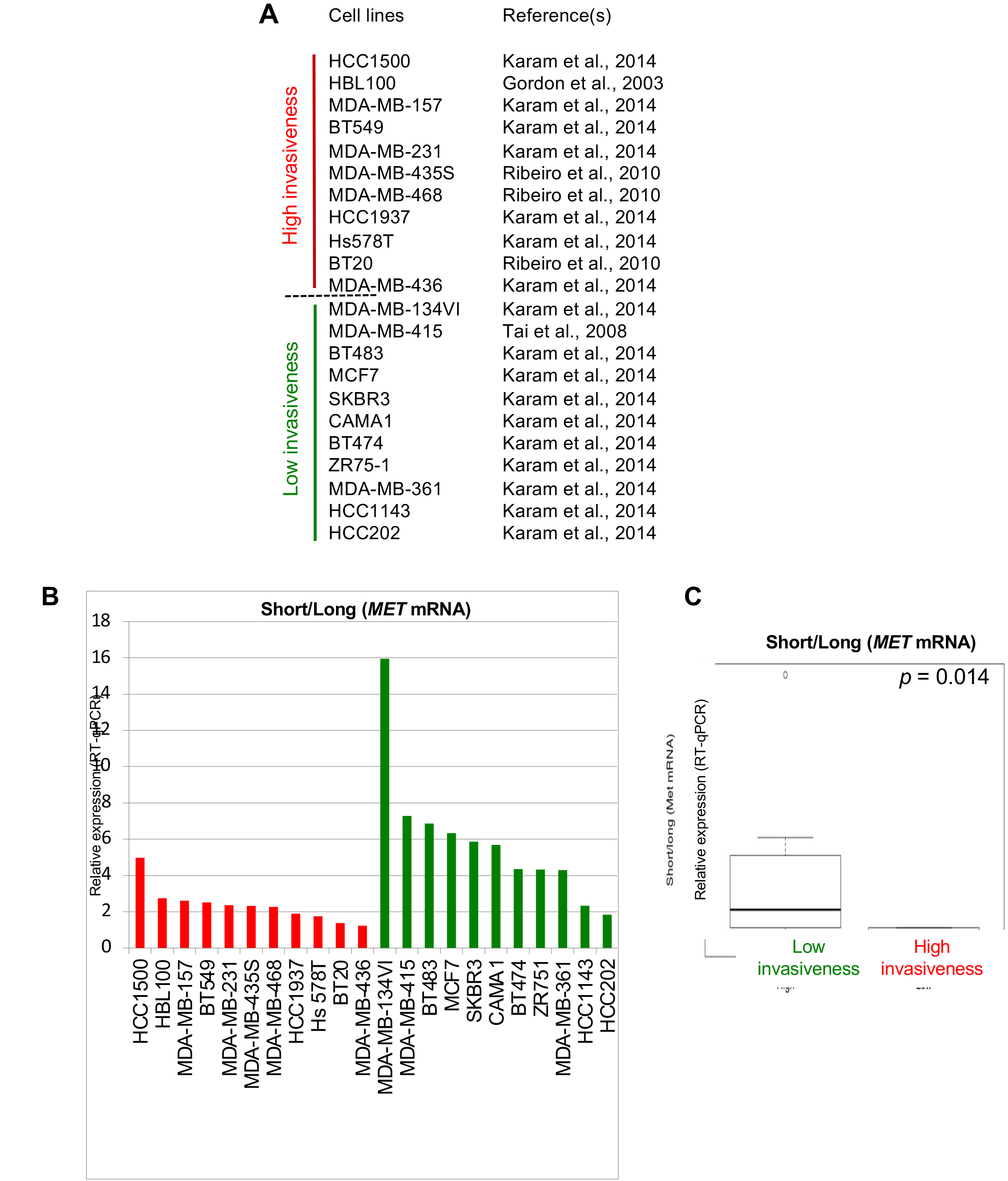
Ratio of short/long MET mRNA isoforms is down-regulated in breast tumor cell lines with high invasive capacities. **A.** List of breast tumor cell lines grouped according to their invasive properties. **B.** RT-qPCR analysis showing the relative expression of short and long MET mRNAs in the indicated cell lines. **C.** Box-and-whisker plot showing the ratio of short/long MET mRNAs.

**Supplementary Figure 2.**
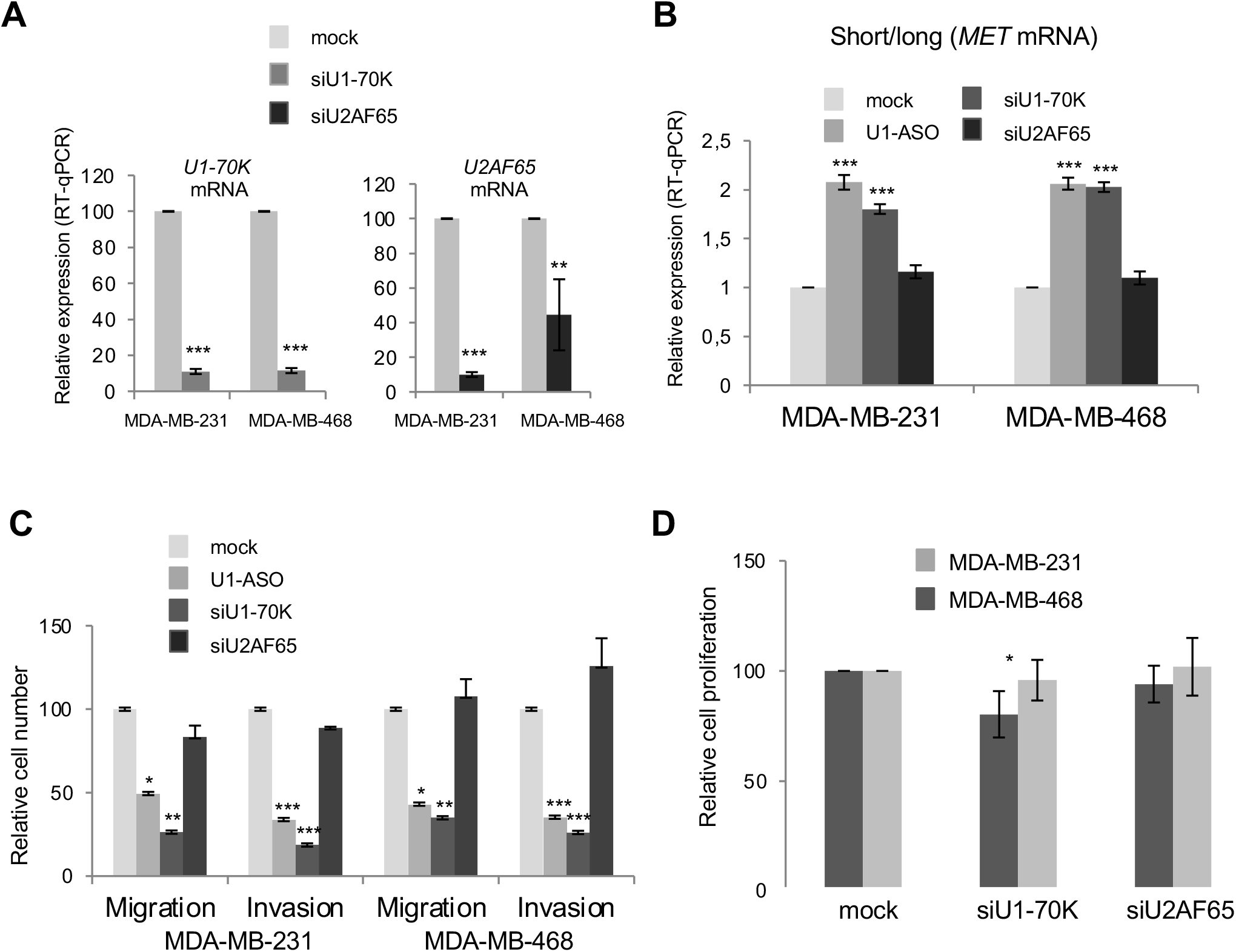
Effect of U1 on in vitro cell migration and invasion is not linked to splicing inhibition. **A.** Efficiency of U1-70K and U2AF65 mRNA depletion by cognate siRNA was analysed by RT-qPCR. U1-70K and U2AF65 expression were normalized to an endogenous control (TBP). Data are presented as the mean ± s.d., and differences were assessed with Student’s t-test (**, P<0.01, ***, P<0.001). **B.** RT-qPCR showing the relative expression of short and long MET mRNAs in breast cancer cell lines after transfection with U1-ASO or siRNAs targeting U1-70K (siU1-70K) or U2AF65 (siU2AF65). Data are presented as the mean ± s.d., and differences were assessed with Student’s t-test (***, P<0.001). **C.** Relative number of migrating and invasive human breast cancer cells after transfection with U1-ASO or siRNAs targeting U1-70K (siU1-70K) or U2AF65 (siU2AF65). Data are presented as the mean ± s.d., and differences were assessed with Student’s t-test (*, P<0.05; **, P<0.01; ***, P<0.001). **D.** Relative proliferation of MDA-MB-231 and MDA-MB-468 cells after siU1-70K or siU2AF65 transfection (*, P<0.05).

**Supplementary Figure 3.**
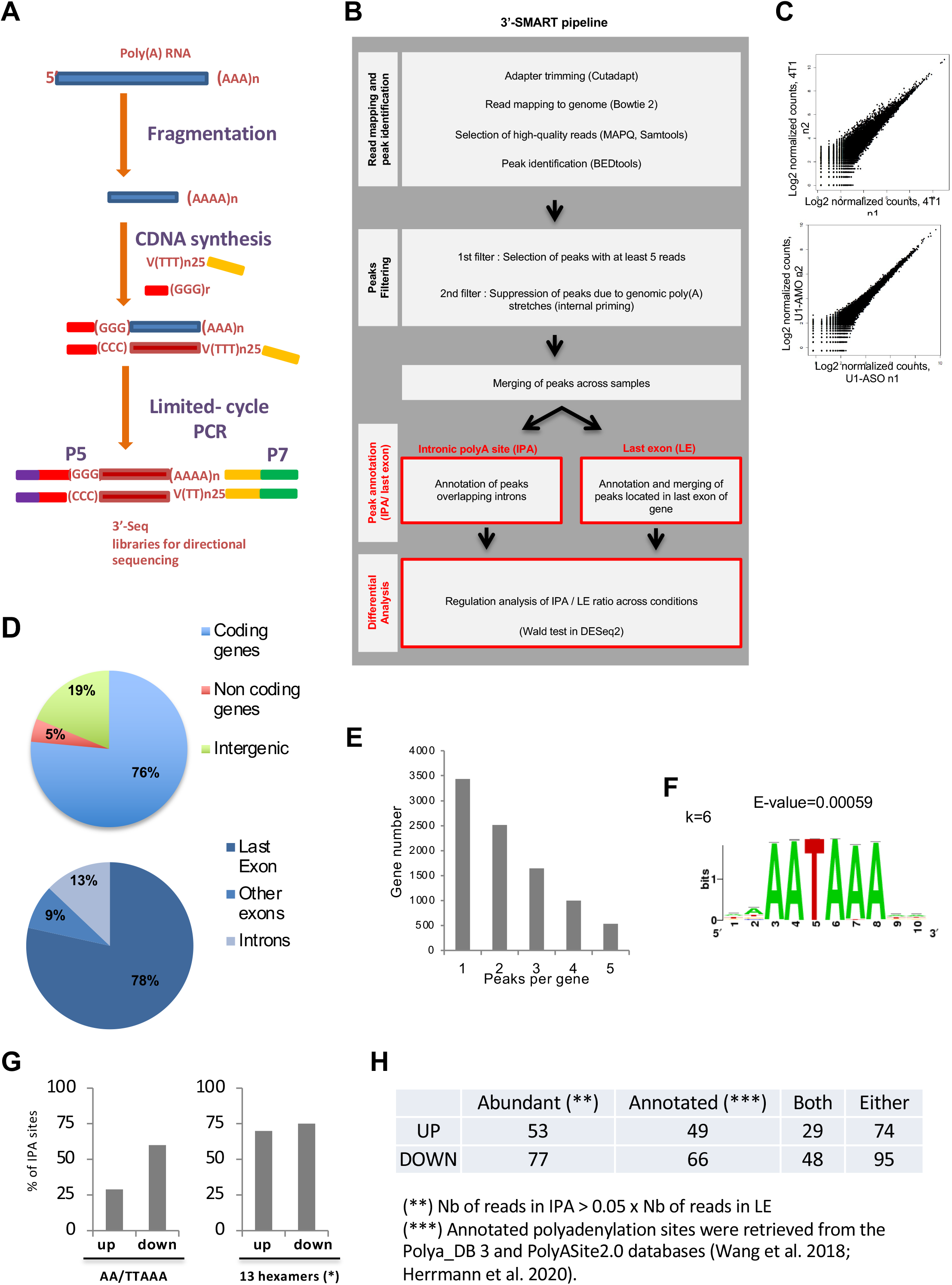
Analysis of IPA regulation using 3’-Seq analysis. **A.** Experimental strategy of 3’-Seq. **B.** Schematic representation of the 3’-SMART pipeline. **C.** Scatter plot of the two biological replicates for each indicated condition. **D.** Pie-chart showing the distribution of 3’-seq reads (4T1 sample) in the mouse genome (upper panel) and within coding genes (bottom panel). **E.** Bar graph showing the number of 3’-seq peaks (pA sites) per gene in 4T1 cells. **F.** The top scoring motif enriched in the peaks identified by 3’-seq in 4T1 cells. **G.** Histogram showing the percentage of down- or up-regulated IPA sites containing a canonical hexameric pA signal, AA/TTAAA or one of the 13 hexamers. **H.** Table showing the percentage of abundant or annotated down- or up-regulated IPA sites.

**Supplementary Figure 4.**
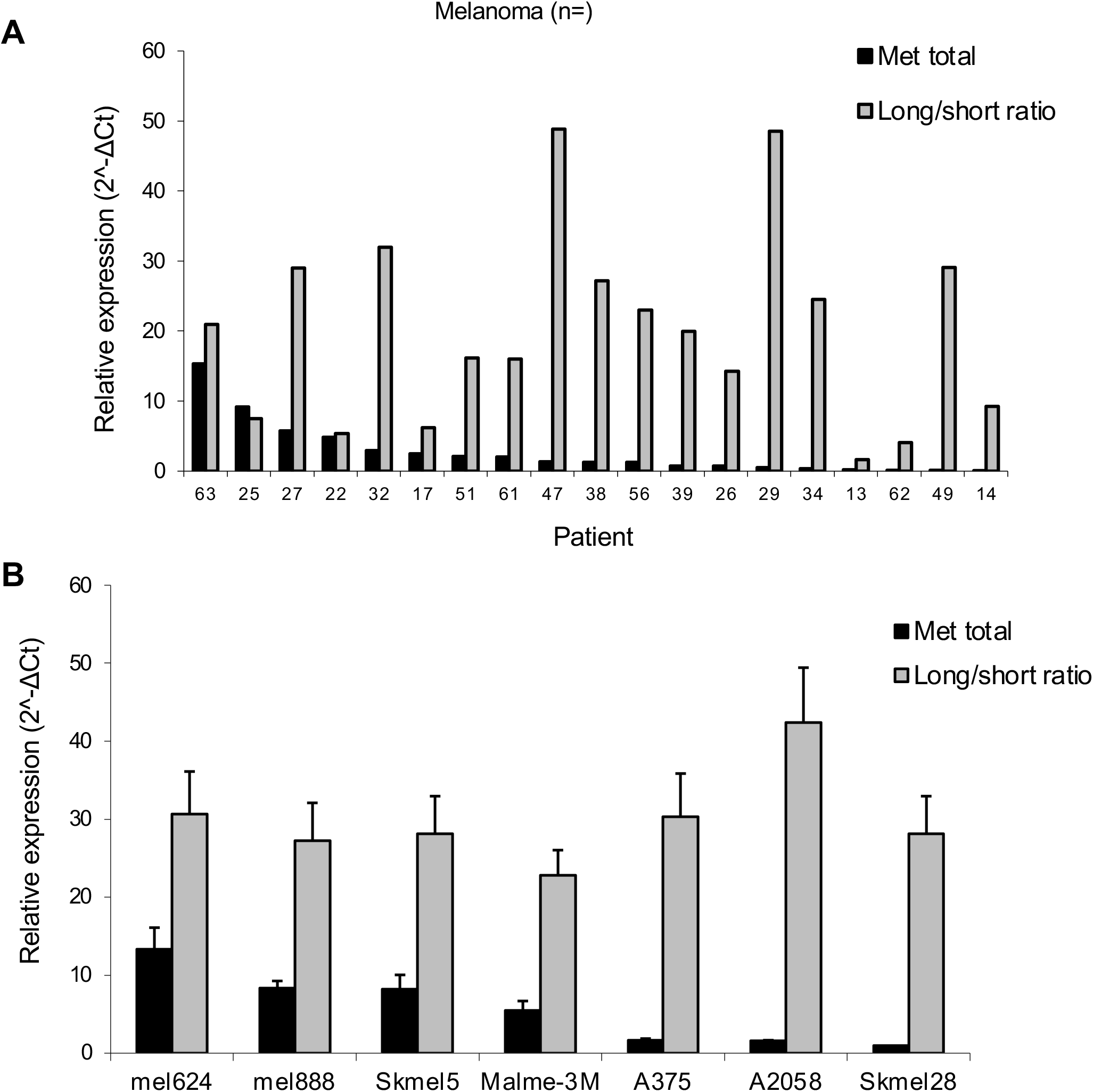
RT-qPCR analysis showing the relative expression of short and long MET mRNAs, as well as the expression of the total MET mRNA (short+long) in 19 melanoma biopsies **(A)** and in the indicated BRAF (V600E) melanoma cell lines **(B)**.

## References

1. Millevoi, S. & Vagner, S. Molecular mechanisms of eukaryotic pre-mRNA 3′ end processing regulation. Nucleic Acids Research 38, 2757–2774 (2010).

2. Tian, B. & Manley, J. L. Alternative polyadenylation of mRNA precursors. Nat Rev Mol Cell Biol 18, 18–30 (2017).

3. Mitschka, S. & Mayr, C. Context-specific regulation and function of mRNA alternative polyadenylation. Nat Rev Mol Cell Biol 23, 779–796 (2022).

4. Berg, M. G. et al. U1 snRNP Determines mRNA Length and Regulates Isoform Expression. Cell 150, 53–64 (2012).

5. Gunderson, S. I., Polycarpou-Schwarz, M. & Mattaj, I. W. U1 snRNP Inhibits Pre-mRNA Polyadenylation through a Direct Interaction between U1 70K and Poly(A) Polymerase. Molecular Cell 1, 255–264 (1998).

6. Kaida, D. et al. U1 snRNP protects pre-mRNAs from premature cleavage and polyadenylation. Nature 468, 664–668 (2010).

7. Vagner, S., Rüegsegger, U., Gunderson, S. I., Keller, W. & Mattaj, I. W. Position-dependent inhibition of the cleavage step of pre-mRNA 3′-end processing by U1 snRNP. RNA 6, 178–188 (2000).

8. Vorlová, S. et al. Induction of Antagonistic Soluble Decoy Receptor Tyrosine Kinases by Intronic PolyA Activation. Molecular Cell 43, 927–939 (2011).

9. Elkon, R. et al. E2F mediates enhanced alternative polyadenylation in proliferation. Genome Biol 13, R59 (2012).

10. Masamha, C. P. et al. CFIm25 links alternative polyadenylation to glioblastoma tumour suppression. Nature 510, 412–416 (2014).

11. Mayr, C. & Bartel, D. P. Widespread Shortening of 30UTRs by Alternative Cleavage and Polyadenylation Activates Oncogenes in Cancer Cells.

12. Chan, J. J. et al. Pan-cancer pervasive upregulation of 3′ UTR splicing drives tumourigenesis. Nat Cell Biol 24, 928–939 (2022).

13. Ichinose, J. et al. Alternative polyadenylation is associated with lower expression of PABPN1 and poor prognosis in non-small cell lung cancer. Cancer Science 105, 1135– 1141 (2014).

14. Lai, D.-P. et al. Genome-wide profiling of polyadenylation sites reveals a link between selective polyadenylation and cancer metastasis. Human Molecular Genetics 24, 3410– 3417 (2015).

15. Zhao, Z. et al. Comprehensive characterization of somatic variants associated with intronic polyadenylation in human cancers. Nucleic Acids Research 49, 10369–10381 (2021).

16. Zhao, Z. et al. Cancer-associated dynamics and potential regulators of intronic polyadenylation revealed by IPAFinder using standard RNA-seq data. Genome Res. 31, 2095–2106 (2021).

17. Singh, P. et al. Global Changes in Processing of mRNA 3′ Untranslated Regions Characterize Clinically Distinct Cancer Subtypes. Cancer Research 69, 9422–9430 (2009).

18. Xia, Z. et al. Dynamic analyses of alternative polyadenylation from RNA-seq reveal a 3′-UTR landscape across seven tumour types. Nat Commun 5, 5274 (2014).

19. Dutertre, M. et al. A recently evolved class of alternative 3′-terminal exons involved in cell cycle regulation by topoisomerase inhibitors. Nat Commun 5, 3395 (2014).

20. Tien, J. F. et al. CDK12 regulates alternative last exon mRNA splicing and promotes breast cancer cell invasion. Nucleic Acids Research 45, 6698–6716 (2017).

21. Oh, J.-M. et al. U1 snRNP regulates cancer cell migration and invasion in vitro. Nat Commun 11, 1 (2020).

22. Ma, P. C. MET receptor juxtamembrane exon 14 alternative spliced variant: Novel cancer genomic predictive biomarker. Cancer Discov 5, 802–805 (2015).

23. Arnold, L., Enders, J. & Thomas, S. M. Activated HGF-c-Met Axis in Head and Neck Cancer. Cancers (Basel) 9, (2017).

24. Marona, P. et al. C-Met as a Key Factor Responsible for Sustaining Undifferentiated Phenotype and Therapy Resistance in Renal Carcinomas. Cells 8, (2019).

25. Szturz, P. et al. Prognostic value of c-MET in head and neck cancer: A systematic review and meta-analysis of aggregate data. Oral Oncol. 74, 68–76 (2017).

26. Zhang, H., Feng, Q., Chen, W.-D. & Wang, Y.-D. HGF/c-MET: A Promising Therapeutic Target in the Digestive System Cancers. Int J Mol Sci 19, (2018).

27. Engelman, J. A. et al. MET amplification leads to gefitinib resistance in lung cancer by activating ERBB3 signaling. Science 316, 1039–1043 (2007).

28. Kubo, T. et al. MET gene amplification or EGFR mutation activate MET in lung cancers untreated with EGFR tyrosine kinase inhibitors. Int. J. Cancer 124, 1778–1784 (2009).

29. Shattuck, D. L., Miller, J. K., Carraway, K. L. & Sweeney, C. Met receptor contributes to trastuzumab resistance of Her2-overexpressing breast cancer cells. Cancer Res. 68, 1471–1477 (2008).

30. Straussman, R. et al. Tumour micro-environment elicits innate resistance to RAF inhibitors through HGF secretion. Nature 487, 500–504 (2012).

31. Karamouzis, M. V., Konstantinopoulos, P. A. & Papavassiliou, A. G. Targeting MET as a strategy to overcome crosstalk-related resistance to EGFR inhibitors. Lancet Oncol. 10, 709–717 (2009).

32. Xu, H. et al. Dual blockade of EGFR and c-Met abrogates redundant signaling and proliferation in head and neck carcinoma cells. Clin. Cancer Res. 17, 4425–4438 (2011).

33. Peruzzi, B. & Bottaro, D. P. Targeting the c-Met signaling pathway in cancer. Clin. Cancer Res. 12, 3657–3660 (2006).

34. Steinway, S. N., Dang, H., You, H., Rountree, C. B. & Ding, W. The EGFR/ErbB3 Pathway Acts as a Compensatory Survival Mechanism upon c-Met Inhibition in Human c-Met+ Hepatocellular Carcinoma. PLoS ONE 10, e0128159 (2015).

35. Wu, Y.-L. et al. Does c-Met remain a rational target for therapy in patients with EGFR TKI-resistant non-small cell lung cancer? Cancer Treat. Rev. 61, 70–81 (2017).

36. Drilon, A., Cappuzzo, F., Ou, S.-H. I. & Camidge, D. R. Targeting MET in Lung Cancer: Will Expectations Finally Be MET? J Thorac Oncol 12, 15–26 (2017).

37. Pasquini, G. & Giaccone, G. C-MET inhibitors for advanced non-small cell lung cancer. Expert Opin Investig Drugs 27, 363–375 (2018).

38. Tiran, Z. et al. A Novel Recombinant Soluble Splice Variant of Met Is a Potent Antagonist of the Hepatocyte Growth Factor/Scatter Factor-Met Pathway. Clinical Cancer Research 14, 4612–4621 (2008).

39. Decorsière, A., Cayrel, A., Vagner, S. & Millevoi, S. Essential role for the interaction between hnRNP H/F and a G quadruplex in maintaining p53 pre-mRNA 3′-end processing and function during DNA damage. Genes Dev. 25, 220–225 (2011).

40. Wang, R. & Tian, B. APAlyzer: a bioinformatics package for analysis of alternative polyadenylation isoforms. Bioinformatics 36, 3907–3909 (2020).

41. Herrmann, C. J. et al. PolyASite 2.0: a consolidated atlas of polyadenylation sites from 3′ end sequencing. Nucleic Acids Research gkz918 (2019) doi:10.1093/nar/gkz918.

42. Legendre, M. & Gautheret, D. Sequence determinants in human polyadenylation site selection. BMC Genomics 4, 7 (2003).

43. Seo, M. et al. RNAi-based functional selection identifies novel cell migration determinants dependent on PI3K and AKT pathways. Nat Commun 5, 5217 (2014).

44. Lezcano, C. et al. Evaluation of stromal HGF immunoreactivity as a biomarker for melanoma response to RAF inhibitors. Modern Pathology 27, 1193–1202 (2014).

45. Morris, A. R. et al. Alternative Cleavage and Polyadenylation during Colorectal Cancer Development. Clinical Cancer Research 18, 5256–5266 (2012).

46. Hall, J. Future directions for medicinal chemistry in the field of oligonucleotide therapeutics. RNA 29, 423–433 (2023)

