## Supplementary material for "MET functions in tumour progression and therapy resistance are repressed by intronic polyadenylation": Table S1

**Table S1. Characteristics of the 416 primary breast tumors**

|  | Number of patients (%) | Number of relapse (%) | *P-value ^a^* |
| --- | --- | --- | --- |
| *Total* | 416 (100) |  |  |
| *Age*  ≤50  >50 | 89 (21.4)  327 (78.6) | 33 (37.1)  129 (39.4) | 0.70 (NS) |
| *SBR histological grade* ^b, c^  I  II  III | 54 (13.3)  208 (51.1)  145 (35.6) | 10 (18.5)  83 (39.9)  65 (44.8) | **0.0017** |
| *Lymph node status* ^d^  0  1-3  >3 | 112 (27.2)  216 (52.4)  84 (20.4) | 33 (29.5)  74 (34.3)  53 (63.1) | **0.00000038** |
| *Macroscopic tumor size* ^e^  ≤25mm  >25mm | 206 (50.5)  202 (49.5) | 60 (29.1)  101 (50) | **0.0000051** |
| *ERα status*  Negative  Positive | 111 (26.7)  305 (73.3) | 44 (39.6)  118 (38.7) | 0.27 (NS) |
| *PR status*  Negative  Positive | 176 (42.3)  240 (57.7) | 75 (42.6)  87 (36.3) | **0.049** |
| *ERBB2 status*  Negative  Positive | 330 (79.3)  86 (20.7) | 126 (38.2)  36 (41.9) | 0.44 (NS) |
| *Molecular subtypes*  RH- ERBB2-  RH- ERBB2+  RH+ ERBB2-  RH+ ERBB2+ | 66 (15.9)  40 (9.6)  264 (63.5)  46 (11.1) | 24 (36.4)  19 (47.5)  102 (38.6)  17 (37.0) | 0.29 (NS) |
| *PIK3CA mutation status* ^f^  wild type  mutated | 279 (67.4)  135 (32.6) | 117 (41.9)  44 (32.6) | **0.032** |

a Log-rank test. NS : not significant

b Scarff Bloom Richardson classification.

c Information available for 407 patients.

d Information available for 412 patients.

e Information available for 408 patients.

f Information available for 414 patients.
