## Supplementary material for "MET functions in tumour progression and therapy resistance are repressed by intronic polyadenylation": Table S3

**Table S3: Characteristics of the 78 head and neck squamous cell carcinomas**

|  | Number of patients (%) | Number of progression (%) | *P-value ^a^* |
| --- | --- | --- | --- |
| *Total* | 78 (100) | 65 (83.3) |  |
| *Age*  ≤62  >62 | 43 (55.1)  35 (44.9) | 40 (93.0)  25 (71.4) | **0.041** |
| *Sexe*  Male  Female | 63 (80.8)  15 (19.2) | 55 (87.3)  10 (66.7) | 0.20 (NS) |
| *Tobacco* ^b^  No  Yes | 12 (15.8)  64 (84.2) | 6 (50.0)  57 (89.1) | **0.039** |
| *Alcool* ^c^  No  Yes | 25 (40.3)  37 (59.7) | 18 (72.0)  33 (89.2) | 0.33 (NS) |
| *HPV*  Negative  Positive | 67 (85.9)  11 (14.1) | 60 (89.6)  5 (45.5) | **0.0015** |
| *Stage*  I  II  III  IV | 3 (3.8)  9 (11.5)  27 (34.6)  39 (50.0) | 3 (100)  7 (77.8)  21 (77.8)  34 (87.2) | 0.91 (NS) |
| *Tumor location*  Oral cavity  Oropharynx  Larynx  Hypopharynx  Other  Several  Unknown | 16 (20.5)  33 (42.3)  7 (9.0)  6 (7.7)  5 (6.4)  8 (10.3)  3 (3.8) | 15 (93.8)  26 (78.8)  4 (57.1)  6 (100)  5 (100)  7 (87.5)  2 (66.7) | 0.094 (NS) |

a Log-rank test. NS : not significant

b Information available for 76 patients.

c Information available for 62 patients.
